# Neuronal Decoding of Temperature Signals in *Caenorhabditis elegans*

**DOI:** 10.1101/2025.01.27.634994

**Authors:** Abhilasha Batra, Rati Sharma

## Abstract

Neural processing in animals facilitate sensory adaptation by eliciting appropriate responses to changing environmental stimuli. Although sensory adaptation is well recognized, the specific role of individual neurons in these adaptive mechanisms remains poorly understood from a theoretical standpoint. Specifically, it is unclear how single neurons influence the processing of sensory information and modulate their responses to environmental changes. In *Caenorhabditis elegans* (*C. elegans*), the primary AFD thermosensory neurons respond to external thermal stimuli incorporating experience-dependent temperature response thresholds. While extensive experimental studies have highlighted the functional features of the AFD neurons, mathematical understanding of their responses is still not well established. In this study, we develop a theoretical framework that captures how a single pair of AFD neurons encodes sensory information and regulates their responses to thermal warming signals, with a focus on how prior environmental experiences influence these responses. Through this framework we find that the interaction between two key entities *i*.*e*. cGMP and calcium, plays a crucial role in processing sensory information and fine-tuning neuronal responses to thermal changes. In addition to reproducing experimentally known trends, our results show that growth conditions such as cultivation temperature of the worms significantly shape the neurons’ functional properties, including their operating range, activation thresholds and response start time and duration. The findings in this work, therefore, enhance the overall understanding of how environmental experiences by single neuron shape sensory systems and provide a deeper connection between experimental data and quantitative models.

## INTRODUCTION

Timely and appropriate response to external environmental stimuli is essential to an organism’s perception of its surroundings and subsequent survival. In animals, this perception and response is facilitated by the activation of specific neurons and signal transduction through the neuronal network. Therefore, understanding neuronal signaling and response is key to decoding various environmental stimuli at the cellular level. To this end, in this work, we specifically focus on neuron level decoding of temperature fluctuations in the environment through studies carried out in the model organism, the nematode (worm) *Caenorhabditis elegans*.

As ectotherms, these nematodes quickly adjust their body temperature to match that of their environment [1]. This rapid equilibration with the environment underscores their importance as model systems for studying thermosensory transduction in neurons. In laboratory experiments, the effect of temperature on the behavior of *C. elegans* primarily emerges through their temperature dependent migration, known as thermotaxis [2]. Although this thermotactic behavior results from the proper functioning of the entire neuronal network, previous studies have shown AFD neurons to have a primary role in thermosensation. In fact, a severe disruption in thermotaxis has been observed in worms that have a pair of AFD sensory neurons ablated in them [3, 4]. This then highlights the importance of recognizing that a comprehensive understanding of an animal’s nervous system requires a detailed exploration of the specific roles played by individual sensory neurons, which are crucial for overall sensory processing.

In light of this finding, several experimental studies on *C. elegans* have focused on understanding the response of AFD sensory neurons to the external thermal stimuli by performing calcium imaging on them [5–11]. This approach of monitoring calcium transients takes advantage of the fact that an increase in the activity of the neuron is often characterized by an influx of calcium ions in it [5, 12–14]. A few recent studies have also focused on the spatiotemporal regulation of the second messenger, the molecule cyclic guanosine monophosphate (cGMP), in the neurons. Specifically for AFD neurons, different FRET- and GFP-based cGMP sensors have been applied to visualize their response under external thermal stimuli [15, 16].

Although experimental research has enabled direct monitoring of the dynamical activity of AFD neurons, mathematical models that can capture these response dynamics and offer mechanistic insights are still missing [17, 18]. In the past, modeling studies for other sensory neurons combined with validation through experiments have been effective in representing and capturing key aspects of neuronal dynamics [19–22], thus highlighting the potential of similar approaches for AFD neurons. Therefore, developing a systems-level mathematical framework for the dynamics of signaling entities within the AFD neurons can not only provide insights about sensory information transduction but also predict responses under varying thermal stimulus conditions. Specifically, by investigating the interplay between cGMP and calcium within these neurons, such models can elucidate how thermal signals are encoded and how prior growth conditions shape neuronal properties. Hence, in this study, we attempt to bridge this gap and enhance our understanding of the dynamic relationship between environmental stimuli and neural adaptation, offering a comprehensive perspective on thermosensory processing in *C. elegans*.

In this work, we develop a simplified model of the primary thermotransduction pathway found in the AFD neurons to understand its signal-response dynamics. This model aims to be a valuable theoretical framework that can be used to predict and interpret the complex response dynamics of neurons. The main analysis of the model explores how changes in the cGMP signaling cascade, the main signaling pathway within AFD neurons, influence the dynamics of the concentrations of two important entities, *i*.*e*., cGMP and calcium, in response to a linear thermal warming signal. By focusing on the rate of change of the levels of cGMP and calcium in the neuron, the model also sheds light on how these components contribute to thermal response adaptation in *C. elegans*. Various modeling studies in the past on different neurons have shown that environmental signals can be encoded in multiple ways, including the absolute signal levels, the signal derivative, the signal’s fold change or even the second derivative, among others [23–25]. Building on these insights, we use our mechanistic model of AFD neurons to first investigate whether these neurons encode signals as absolute levels or as temporal changes, with an objective to understand the principles that drive their adaptive response to temperature shifts.

Another important property of thermosensory AFD neurons that has been observed experimentally is that the temperature threshold at which activity is initiated in these neurons depends on the past growth temperature of the worms [10, 26]. Therefore, neurons when given thermal stimuli show changes in their intracellular calcium levels only near a certain response threshold temperature, *T* ^*^, which in turn, is regulated by the growth/cultivation temperature, *T*_*C*_. Remarkably, single-cell studies confirm that AFD neurons retain this temperature information independently of inputs from other cells [27], thus highlighting their capacity to acquire, retain and reset the *T*_*C*_ dependent information. However, to the best of our knowledge, no mathematical model has yet been developed to quantitatively analyze this memory-like behavior and the relationship between *T* ^*^ and *T*_*C*_ across a wide range of thermal signal. Our model, therefore, aims to address this by examining how *T* ^*^ depends on *T*_*C*_ for linear thermal warming ramp as the signal. Additionally, it provides specific temperature and temporal response thresholds of AFD neurons within the physiological temperature range (15°C to 25°C) for *C. elegans*. These findings from the analysis of our model capture the memory retaining feature of the neurons and align really well with the experimental study by Kobayashi *et al*. [27]. Moreover, the model can also be extended to incorporate the reset mechanism, another defining feature of the AFD neuron, thus providing a flexible framework to probe both memory retain and reset properties of the neurons.

Next, the potential of the model is further explored where we predict how the response dynamics of AFD neurons correlate with external thermosensory inputs. Using our model, we examine in detail how changes in the rate of signal upstep influence neural adaptation dynamics and the mathematical decay forms that characterize this adaptation. In addition, the model also effectively captures the high sensitivity of AFD neurons to differential temperature changes in the signal even on the order of 0.1°C. After validating the model against experimental calcium levels reported in a recent study by Huang *et al*. [28], further analysis of the model allows for prediction of response properties such as operating range, relaxation phase, onset time and response time of the neuron. A detailed comparison of these response properties across a range of growth/cultivation temperatures helps in understanding the rate of temperature-evoked activity in the AFD neurons of *C. elegans*. Overall then, the model developed in this work enhances the quantitative understanding of the response evoked in the AFD neuron responses and broadens the scope for investigating mechanisms of neuronal adaptation, particularly of thermosensation in *C. elegans*.

## RESULTS

### A simplified model of the thermosensory transduction pathway in *C. elegans*

The bilateral pair of bipolar AFD sensory neurons reside in the head of *C. elegans*, with its dendrites extending to the tip of the nose and axons entering into the nerve ring [29, 30]. The AFD neurons are unique in their ability to detect and respond to even tiny thermal fluctuations. This extraordinary thermosensitivity and broad dynamic range of AFD neurons is provided by the activation of an intracellular signaling pathway that involves a complex cGMP dependent thermotransduction cascade [31, 32], shown in Fig. 1a. This pathway begins with the activation of thermosensitive receptor guanylyl cyclases (rGCs), which detect temperature changes and initiate a signaling cascade. The signal transmission in AFD neurons mainly rely on cGMP as a second messenger, which activate downstream ion channels thus leading to the calcium response in the neuron. The topology of the cGMP-dependent cascade that includes all these steps is depicted in Fig. 1b and is analogous to the signal transduction and adaptation pathway deduced from various experimental studies [33].

**Figure 1.**
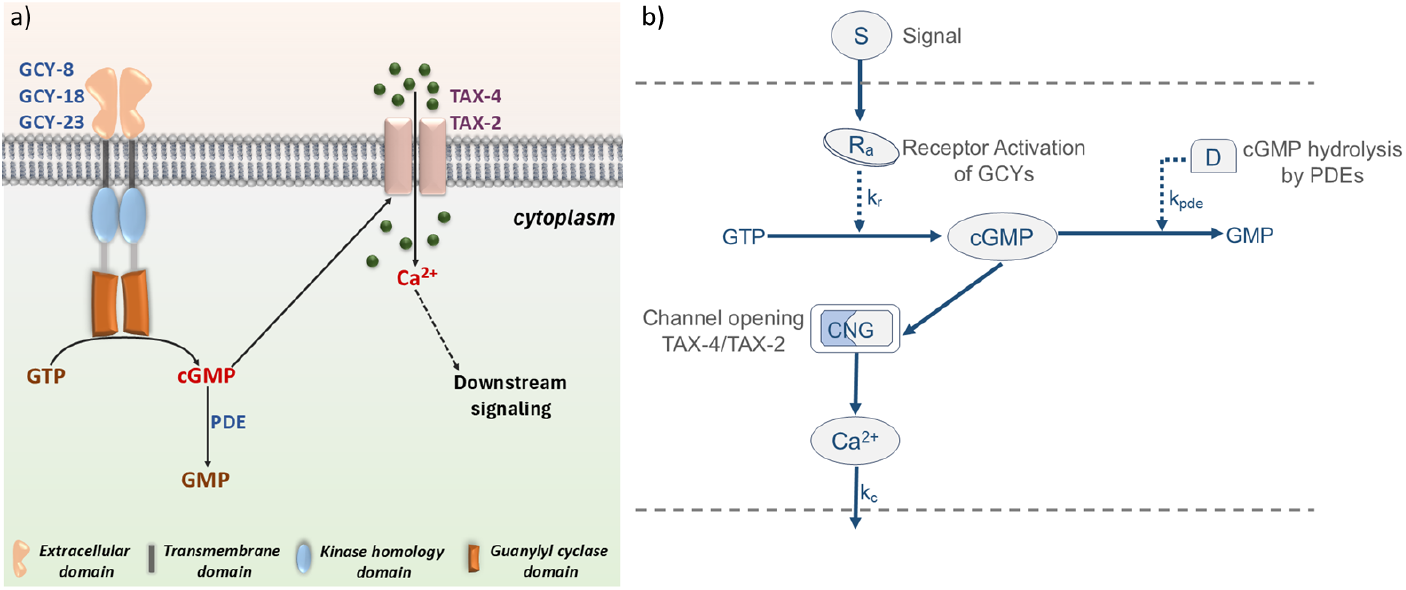
Illustration of thermosensory transduction pathway in the AFD neurons of *C. elegans* and the analogous model: **a)** Upon warming, the receptor guanylyl cyclases (rGCs) GCY-8, GCY-18 and GCY-23 of the AFD neurons get activated to increase the intracellular concentration of second messenger cGMP through their catalytic activity. cGMP gates TAX-4/TAX-2 encoded cation channels that allow the influx of Ca^2+^ ions. Phosphodiesterase (PDEs) mediate cGMP hydrolysis to terminate the signaling in AFD neurons. **b)** An analogous circuit model that recapitulates the cGMP-dependent cascade of the AFD neuron. Each arrow depicts the corresponding reactions between the different components of the model and symbols next to arrows mark the relevant constant in the model’s equations. Note that dashed arrows represent enzymatic reactions.

To understand how efficiently thermosensory AFD neurons autonomously decode and respond to the signals, we consider the biochemical signaling interactions within the cGMP signaling cascade and develop a parsimonious model using a system of time-dependent differential equations. To develop this model, we first propose the governing kinetic rate equations for each part of the cGMP signaling cascade illustrated in Fig. 1b, followed by a discussion of the specifics of each step.

#### Activation of receptor-type transmembrane guanylyl cyclases (rGCs) in response to environmental temperature stimuli

Out of 27 genome encoded rGCs in *C. elegans*, three of them, GCY-8, GCY-18 and GCY-23 are expressed exclusively at the sensory endings of the AFD neuron. These rGCs collectively function as homodimers in the neuron and act as a receptor that gets activated by external temperature stimuli. The extracellular domain of these rGCs senses temperature changes, transitioning them into their active state, *R*_*a*_. In our model, we consider

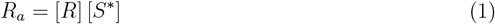

where [*R*] is the initial concentration of rGCs present on the AFD neuron and [*S*^*^] is the level of thermal signal encoded at the sensory endings of neurons by those rGCs. Since linear thermal warming has been the most prevalently used stimulus in experiments that measure the thermosensitivity of AFD neurons [10, 34, 35], to best capture the signal dynamics, we choose to model the signal input, *S*, as a sigmoidal function of temperature. Therefore, *S* is given by

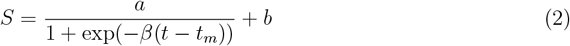

which accurately represents a linear thermal ramp. The numerical value of *a* scales the range of the sigmoidal function, in this case the temperature range and the constant *b* sets the offset temperature in the function. Further, the parameter *β* in the above equation (Eq. 2) determines the rate of transition between temperatures *b* and *a* + *b*, whereas *t*_*m*_ is the characteristic time point that represents the midpoint of the sigmoidal transition. The signal considered in our study matches with experimental studies and varies approximately between 14°C and 24°C with a slope of 0.02°C/*s*. The fit of the time-dependent signal function, *S*, to the experimental data is shown in Fig. S1.

#### cGMP production and degradation

In the neuron, the catalytic activity of the rGC’s kinase domain enables conversion of GTP to its cyclic form, cGMP. Since cGMP synthesis is regulated by an enzyme-activated reaction, it can be modeled through Michaelis-Menten kinetics. The biochemical reaction itself can be written as,

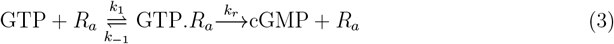

Therefore, the rate of formation of cGMP through enzymatic activity of rGCs is given by

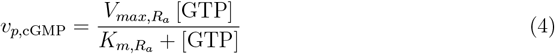

where, 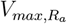 represents the maximal rate of cGMP production. Assuming that the presence of GTP is abundant, 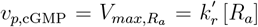,such that 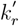 is the product of the rate constant for the production of cGMP, *k*_*r*_, and 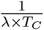,where *T*_*C*_ is the growth/cultivation temperature of the worms and *λ* is the scaling factor.

Further, various experimental studies suggest that together with the rGCs, multiple isoforms of cyclic nucleotide phosphodiesterases (PDEs) also play a critical role in regulating temperature-dependent cGMP levels in AFD neurons [36, 37]. Different isoforms of PDEs (PDE-1, PDE-2, PDE-3 and PDE-5) catalyse hydrolysis of cGMP into GMP to finally terminate the response of the neuron. This reaction can therefore be represented as,

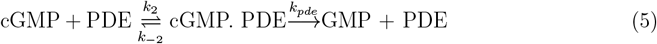

Assuming that intracellular PDEs are present at a constant concentration and have homogeneous catalytic affinity to its substrate cGMP, the rate of cGMP degradation using Michaelis-Menten kinetics is given by,

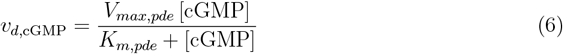

where, *V*_*max,pde*_ represents the maximum rate of cGMP hydrolysis. Previous experiments in mammalian cells have shown that the activity of PDEs increases up to 9-11 times when cGMP binds to its N-terminal domain [38–41]. The mechanism of PDE activation by cGMP can therefore be described as cGMP + PDE_*inact*_ ↔ PDE_*act*_. Consequently, the concentration of active PDE depends on the level of cGMP, because in a Michaelis-Menten kinetics *V*_*max,pde*_ is proportional to the total concentration of active PDEs, which we assume is approximately proportional to the cGMP concentration. This assumption is reasonable if the PDEs levels are high compared to the cGMP levels. Therefore, we describe the maximum cGMP hydrolysis rate *V*_*max,pde*_ as being proportional to cGMP concentration, such that,

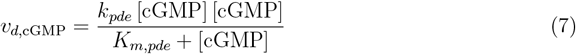

#### Calcium influx and efflux in the AFD neuron

As cGMP levels increase within the AFD neuron, it leads to the opening of the cyclic nucleotide-gated (CNG) channels to transmit the signal. These TAX-4/TAX-2 channels in *C. elegans* contribute to the spatio-temporal dynamics of internal calcium in a manner similar to that of CNG channels in vertebrates [42, 43]. In these organisms, genes *tax-4* and *tax-2* encode the heteromeric CNG-type channel where TAX-4 acts as the *α* subunit and TAX-2 as a modifying *β* subunit [44]. Considering the gating of these channels by cGMP to be cooperative, the cGMP-dependence of channel currents can be described by the Hill equation, where the influx rate is given by

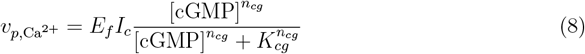

Here, *I*_*c*_ is the circulating current, *n*_*cg*_ is the Hill coefficient, *K*_*cg*_ is the half-saturating concentration of cGMP and *E*_*f*_ is a constant that changes the electrical current unit to concentration changes while taking into account the volume of the AFD neuron. Therefore, *E*_*f*_ is given by

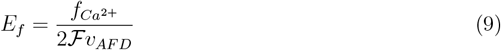

where 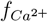 is the fraction of current carried by Ca^2+^ ions, ℱ is the Faraday’s constant, *v*_*AF D*_ is the average volume of the pair of AFD neurons and the factor of 1*/*2 in the above expression accounts for the charge of a calcium ion. The detailed calculation for obtaining the numerical value of *E*_*f*_ for AFD neurons are discussed in the Supporting Information.

As intracellular calcium concentration rises upon the opening of TAX-4/TAX-2 channels, a first-order release through calcium pumps during downstream signaling helps restore the levels to their baseline value, denoted by 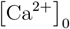,with a removal rate constant *k*_*c*_. Therefore, Ca^2+^ efflux can be finally written as

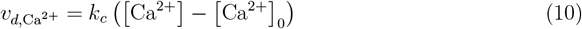

Thus, the dynamics of the cGMP signaling cascade within the AFD neurons, with a focus on the temporal changes of the key signaling components, *i*.*e*., cGMP and Ca^2+^, can be expressed as

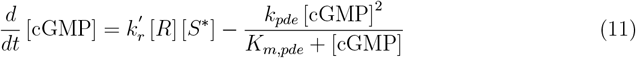

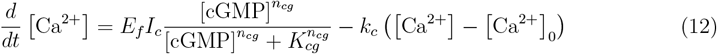

Eqs. 11 and 12 are the key equations that capture the essential intracellular molecular events driving calcium responses within AFD neurons. In all, the model has two key variables and nine parameters, whose numerical values as used in this study and associated references are provided in Table 1. Even though the model, as summarized by Eqs. 11 and 12, seems simple at the outset, the intricacies within each term are quite detailed and complex. However, by focusing on the key molecular players, cGMP and Ca^2+^, this model highlights the dominant dynamics believed to govern the cGMP signaling cascade in response to thermal stimuli. This approach, grounded in both experimental observations [34, 45–48] and theoretical insights, provides a foundation for understanding the interplay between cGMP production and calcium regulation within the thermosensory signaling pathway of AFD neurons. This thus provides a valuable tool for understanding how these signaling components interact and influence AFD neuronal responses to temperature changes. The following sections will provide a detailed analysis of the signal and response as elucidated by this model and also chart out comparisons to and predictions for experiments.

**Table I.** Parameter values used to simulate Eqs. 11 and 12 of the AFD neurons thermotransduction model.

| Parameter | Description | Value | Units | References |
| --- | --- | --- | --- | --- |
| $k_r$ | Rate constant of cGMP synthesis | 3.2 | $\mu s^{-1}$ | Determined from [49] |
| $\lambda$ | Scaling factor in cGMP production | (for $S$) 1 | - | This paper |
| | | (for $dS/dt$) 1 | $s^{-1}$ | This paper |
| $k_{pde}$ | Efflux rate of cGMP catalysed by PDEs | $10^6$ | $s^{-1}$ | This paper |
| $K_{m,pde}$ | Michaelis-Menten constant | 2 | $\mu M$ | [50] |
| $E_f$ | Calcium current to concentration change constant for AFD neurons | $32.33 \times 10^{12}$ | $\mu M C^{-1}$ | Calculated (SI) |
| $I_c$ | Circulating current | 8 | pA | [31] |
| $n_{cg}$ | Hill-coefficient for TAX-4/TAX-2 channel | 0.97 | | [44] |
| $K_{cg}$ | Half saturating concentration of cGMP | 8.4 | $\mu M$ | [44] |
| $k_c$ | Rate constant of calcium removal | 0.08 | $s^{-1}$ | Assumed (based on [24]) |
| Initial values | Description | Value | Units | References |
| $R$ | Initial concentration of rGCs | $\sim 10^6$ | $\mu M$ | This paper |
| cGMP | Initial concentration of cGMP | 0 | $\mu M$ | This paper |
| $Ca^{2+}$ | Initial concentration of $Ca^{2+}$ | 0.1 | $\mu M$ | [51] |

### Adaptive temporal encoding of thermal stimuli in AFD neurons

An animal’s survival and well-being relies heavily on how efficiently it processes signals from its surroundings. To understand how the nervous system of an animal interprets these signals, we must understand how its sensory neurons at the individual level encode and transmit them. Various studies have shown the ability of single neurons to perceive the complex stimuli and perform relevant computations to guide the behavior of *C. elegans* [24, 27, 52]. Therefore, as a first step, in this work, we examine the individual signal encoding properties of AFD neurons, specifically focusing on how they process and respond to gradual and linear increase in temperature.

Certain neurons exhibit characteristic adaptation to prolonged or repeated presentation of stimulus, wherein the response starts to decay after showing an initial onset response to the stimuli [53]. AFD neurons are also known to behave similarly, and therefore, accurately capturing this adaptive feature is essential for any mathematical model aimed at describing neuronal responses. However, different neurons employ different encoding strategies to process the signals they receive, thereby, affecting the response dynamics and adaptation as well. AFD neurons, in particular, are known to regulate thermotaxis and respond to minor temperature changes as well primarily because of their sensitivity to the time derivative of the thermal signal [10, 11]. In light of this, our model explores the sensory-encoding principle by comparing two encoding strategies: one based on absolute signal levels (*S*) and the other on the first derivative (*dS/dt*) of the signal. The analysis of the model considering the encoding by AFD neurons as the absolute signal levels, *S* and the corresponding Ca^2+^ response is shown in the top panel of Fig. 2. These responses show an increase in the level of Ca^2+^ after an upstep in temperature. However, the responses fail to gradually reach the pre-stimulus levels after attaining their peak values. Thus, the possibility that AFD neurons encode the absolute signal levels overlook the important aspect of neural adaptation to thermal stimuli. In contrast, when the neuron encodes or senses the signal as a first derivative of the thermal ramp, represented as *dS/dt* in the model, the model successfully captures the experimental features of cGMP [15, 16] and Ca^2+^ [10, 11] response dynamics, as shown in Fig. S2 and the bottom panel of Fig. 2, respectively. This adaptive thermosensory response of the AFD neuron, depicted in Figs. S2c and 2c, therefore, underscores the important feature of AFD neurons where they compute changes in the temporal activity of temperature signals rather than focusing only on the absolute levels. Thus, by incorporating a first-derivative encoding in our model, we enhance its potential to accurately represent the response dynamics of AFD neurons. Therefore, the final expression for the signal encoded (*S*^*^) at the sensory endings of the receptor in Eq. 1 (plotted in Fig. 2b) is given by,

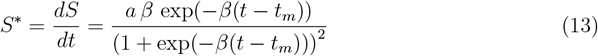

**Figure 2.**
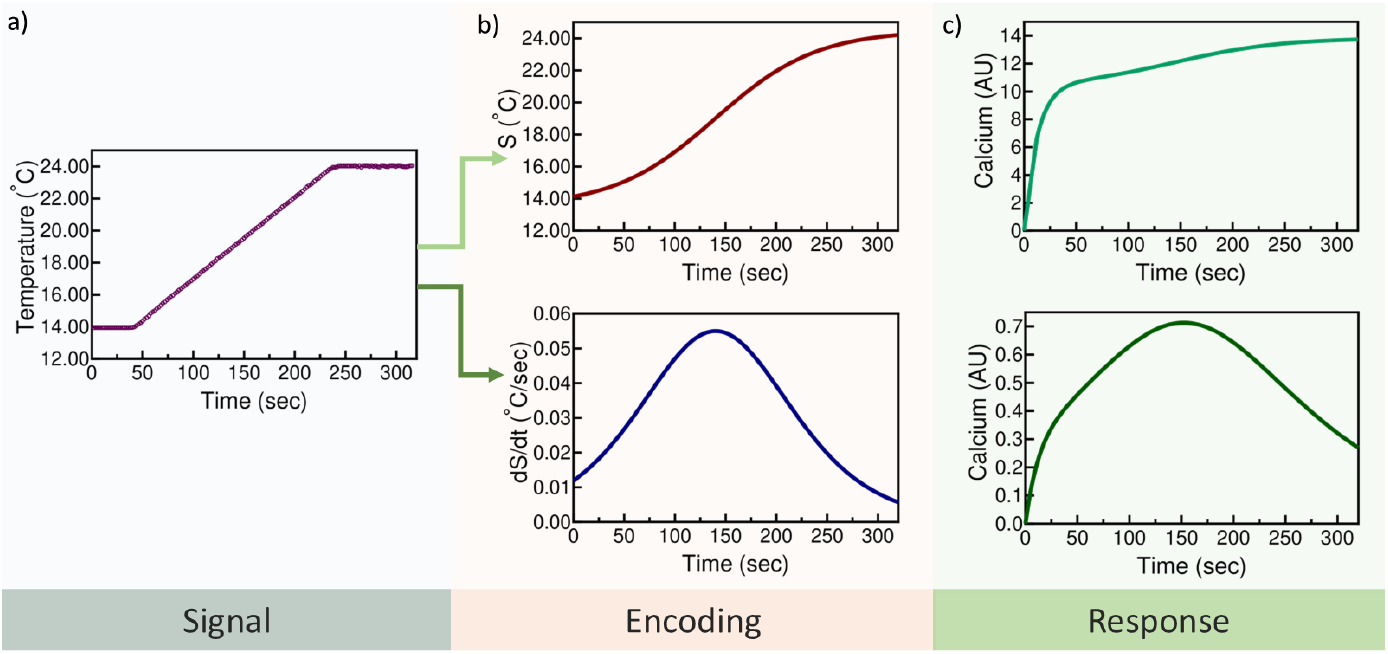
Signal processing in AFD neurons: **a)** Linear thermal warming signal employed in the experimental study [27] on AFD neurons. **b)** *top*: Absolute signal encoding by the neurons with signal, *S*, having mathematical form of a sigmoidal function described by Eq. 2. *below* : Encoding of signal as first derivative, *dS/dt*, (Eq. 13) by the neurons. **c)** Ca^2+^ dynamics observed through the model analysis for encoding of signal as absolute levels *S* (*top*) and *dS/dt* (*below*) at *T*_*C*_ = 20°C.

### Prior temperature experiences are encoded in the response dynamics of AFD neurons

*C. elegans* acclimate to a physiological temperature range of 15°C - 25°C in their environment and are known to perform experience-dependent navigation determined by their growth/cultivation temperature, *T*_*C*_. This thermal memory of *T*_*C*_ is encoded and stored during the development of worms and lasts all through their month-long lifespan [35, 54]. The memorization of prior growth conditions allow the worms to modulate their behavioral response thresholds, enabling them to respond over a broad range of thermal stimuli. Upon experiencing spatial thermal gradients, the worms perform thermotaxis [54, 55]. They move towards colder temperatures, termed negative thermotaxis, when experiencing a temperature *T* above *T*_*C*_ and move towards higher temperatures, termed positive thermotaxis when *T < T*_*C*_. They also display isothermal tracking when the temperature is kept same as *T*_*C*_. Previous genetic studies suggest that the behavioral adaptation of worms in this *T*_*C*_ dependent manner is partly mediated by the signaling cascade of AFD thermosensory neurons [2, 3, 26, 56]. Further, calcium imaging studies on the AFD neurons support this observation as the response to thermal stimulus ramp is initiated only above a certain threshold temperature, *T* ^*^, also called response temperature. This threshold for temperature-evoked activity is determined by the worms’ growth/cultivation temperature and is regulated by intracellular cGMP levels [57]. Since AFD neurons can independently acquire, store and reset the growth temperature information without input from other cells [27], investigating this feature using a mathematical framework specifically for AFD neurons will offer deeper insights, which we next explore in this study.

To begin understanding how *T*_*C*_ regulates and sets the threshold for temperature-evoked activity in AFD neurons, we first examine the relationship between *T*_*C*_ and *T* ^*^. Using the experimental study by Kobayashi *et al*. [27], we first establish that *T* ^*^ varies linearly with *T*_*C*_, *i*.*e*., *T* ^*^ = 0.79 *T*_*C*_ + 3.1 (details are discussed in *Materials and Methods*). Since Eq. 2 in our model represents the time-dependent variation of temperature stimuli, the absolute value of 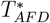 corresponding to a given *T*_*C*_ is applied in it to determine the respective response time, denoted as *t*_*R*_ (Fig. S4a). Previous experimental studies have shown that severing the dendrites of the AFD neuron also destroys the rGCs localized there and thus abolishes their Ca^2+^ response activity [10]. This supports the assumption that memory is stored at the receptors of the neuron and is recalled upon exposure to thermal stimuli. Therefore, we refine our model for the signaling cascade of the AFD neuron by incorporating a time dependent memory recall function into the activated receptors, *R*_*a*_, in Eq. 1, such that,

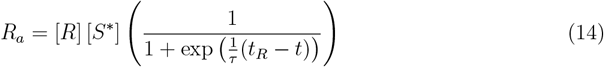

Here,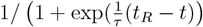 is the memory recall function which is a type of logistic function introduced to temporally modulate the neuronal response with its integration to past growth temperature. The exponential dependence on the time difference, *t*_*R*_ − *t* (where *t*_*R*_ is determined by *T*_*C*_) sets the activation threshold for cGMP production in a manner such that time *t < t*_*R*_ leads to suppressed production of cGMP. On the other hand, the time-point at which *t* approaches *t*_*R*_ marks the activation time point of cGMP production and higher values of *t* leads to maximum activation of cGMP synthesis. The parameter *τ* in the recall function allows fine-tuned control over how quickly the cGMP response is activated, where a smaller value of *τ* results in a rapid shift around *t*_*R*_, while a larger value leads to a gradual increase in the cGMP activation. For the model analysis, *τ* in the recall function has numerical value 1 with *s*^−1^ as the units. Therefore, the differential equation defined for cGMP response dynamics is modified with growth condition dependent memory recall and is expressed as,

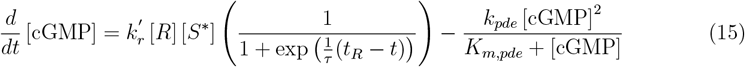

Now, we analyze the model by solving the coupled differential equations, *i*.*e*., Eqs. 15 and 12, to explore how the dynamics of cGMP and Ca^2+^ responses are influenced by *T*_*C*_ dependent memory for the temperature signal described in Eq. 2. We find that the initiation of the AFD neurons’ response, marked by an increase in Ca^2+^ levels, begins at a threshold temperature, *T* ^*^, for each specific *T*_*C*_ and at a corresponding response time, *t*_*R*_. This is shown in Fig. 3a-c. Consistent with observations from several experimental studies [10, 26, 27], this threshold deciding response temperature, *T* ^*^, observed from the model analysis in this study has values close to the respective *T*_*C*_’s. Here, we specifically provide the detailed comparison for the temporal response of both Ca^2+^ (Fig. 3a-c) and cGMP (Fig. S3a-c) at three different *T*_*C*_ values - 17°C, 20°C and 23°C and observe that higher growth temperatures result in the delayed and shifted initiation of the response, along with a faster rate of response adaptation. Having established that the model effectively captures the AFD neuron’s ability to retain temperature memory, we next explore whether the model is sufficiently broad to explain the *T*_*C*_ dependent memory reset, a phenomenon previously observed in temperature shift experimental studies [27]. In order to evaluate the model’s capability for analyzing and replicating the AFD neuron’s temperature memory reset feature, the activated receptor, *R*_*a*_, can be defined by introducing a modified memory recall function as,

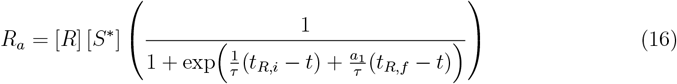

**Figure 3.**
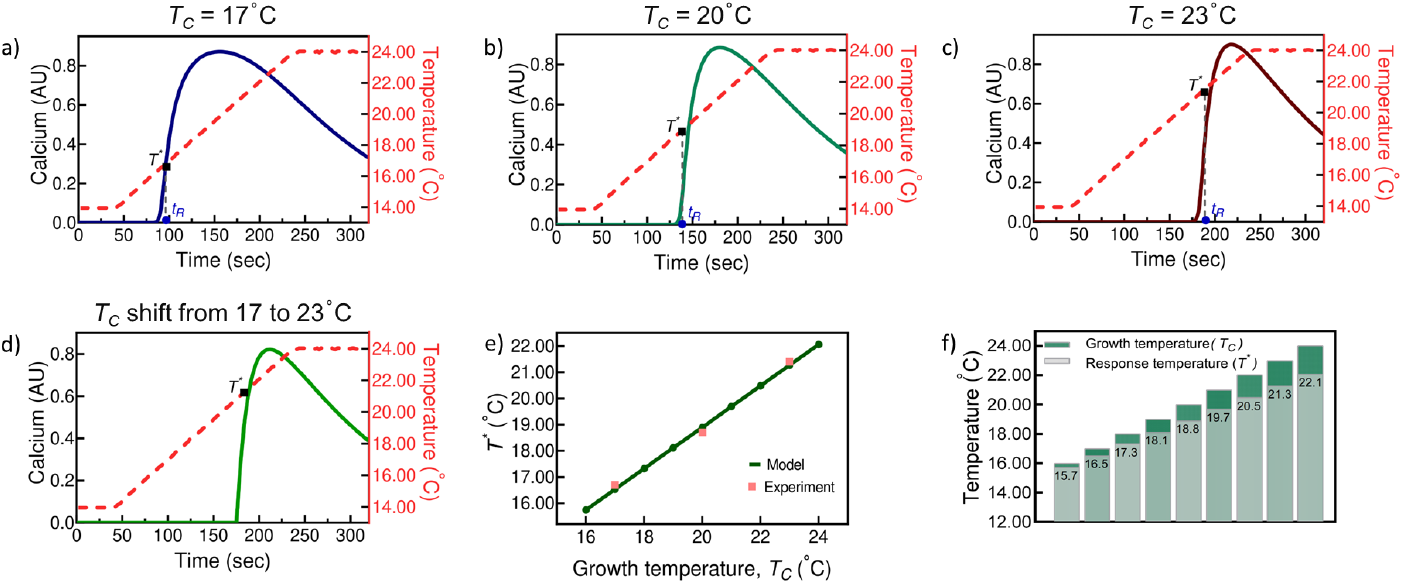
Effect of growth-temperature memory on calcium levels: Ca^2+^ response dynamics in AFD neurons for linear thermal ramp as signal (Eq. 2 and Fig. S1) at **a)** *T*_*C*_ = 17°C, **b)** *T*_*C*_ = 20°C, **c)** *T*_*C*_ = 23°C, **d)** Shift in *T*_*C*_ from 17°C to 23°C. **e)** Comparison between response temperature, *T* ^*^, of the neurons obtained from the model to experimental values observed in [27]. **f)** Change in values of *T* ^*^ from the model analysis for *T*_*C*_ varying in a range of 16°C to 24°C.

Here, the exponential term in the memory recall function depends on the time differences associated with two different *T*_*C*_ conditions. In Eq. 16, *t*_*R,i*_ and *t*_*R,f*_ represent the response times corresponding to temperature transitions, where *t*_*R,i*_ is the time associated with the initial temperature that the neuron experiences, while *t*_*R,f*_ refers to the time associated with the final temperature to which the neuron is shifted. The constant *a*_1_ in Eq. 16 is the scaling factor influencing how strongly the memory reset is affected by the transition to the new temperature. The value *a*_1_ = 10 is optimized for the specific case of memory resetting from a temperature of 17°C to 23°C over a period of more than 6 hours. The results of the model, taking into account the effect of memory reset (from 17°C to 23°C over several hours) on the Ca^2+^ and cGMP levels of the neuron, are shown in Fig. 3d and Fig. S3d, respectively. It can clearly be seen that *T* ^*^ observed due to the temperature memory reset from 17°C to 23°C has a value close to the one observed for the new *T*_*C*_, *i*.*e*., 23°C. This similarity becomes evident on comparing Fig. 3d with Fig. 3c. Further, these results from the analysis of our model with memory reset are found to be in strong agreement with the experimental findings reported in the study by Kobayashi *et al*. [27]. Therefore, the match between response dynamics of neurons measured experimentally to that estimated from the model shows the model’s effectiveness in accurately reproducing the memory retention and reset features of the AFD neurons.

Further, a mathematical model can be particularly valuable in exploring the features of a system that may be challenging to study experimentally. Therefore, using our model (Eqs. 15 and 12) we examine the relationship between the threshold response temperature, *T* ^*^, of the AFD neuron and the cultivation temperature, *T*_*C*_, of the worms across the entire physiological range of 15 − 25°C. From the model we find the *T* ^*^ values to be 16.5, 18.8 and 21.3°C for worms cultivated at 17, 20 and 23°C, respectively. A comparison of the results of the model with the experimental data from Kobayashi *et al*. [27] in Fig. 3e further validates the model’s accuracy and highlights its use in elucidating the neuron’s response properties. Additionally, the change in response time *t*_*R*_ with varying *T*_*C*_ is also deduced from the model and is shown in Fig. S4a. Also, from the study of the model as seen in Fig. 3f, we observe that higher *T*_*C*_ not only leads to correspondingly higher *T* ^*^, but also leads to a wider gap between them. Therefore, the model along with providing good agreement with experimental findings [27], indicates that higher growth temperature conditions significantly shift the response threshold of AFD neurons during linear thermal warming.

### Rate and range of thermal ramp govern the duration of response and adaptation

Having established the relationship between growth temperature and response parameters, we next look at how the rate or steepness of the thermal ramp affects the response activity and adaptation. We do this by simulating the model (Eqs. 15 and 12) for worms with *T*_*C*_ = 20°C but experiencing linear thermal warming at three different rates (details in Table S2). The results are presented in Fig. 4, where Fig. 4a shows the change in Ca^2+^ levels for a thermal stimuli rising gradually with a slope of 0.01°C/*s*. The response dynamics for faster stimulus inputs with a rate of 0.1°C/*s* and for a unit step like temperature change, where the thermal stimulus is shifted from 14°C to 24°C within 10 seconds (with a slope of 1°C/*s*) are shown in Fig. 4b and Fig. 4c, respectively. The corresponding dynamics for the cGMP levels are shown in Fig. S5. Through our model analysis as per Fig. 4 and Fig. S5, we observe that the rate of change in the temperature signal from slow to fast moderately influences the rise dynamics of the neuron’s response to stimuli. As shown in Fig. 4a-c, the time it takes for Ca^2+^ to reach its peak concentration becomes progressively shorter, with the half-rise time shifting by 5.4 seconds to the right as the stimulus rate increases from 0.01 to 1°C/*s*. Next, on comparing the full width at half maximum (FWHM) of the AFD neuron’s response across different signal rates, we found a reduced spread of the Ca^2+^ response with faster rise in temperatures, with the FWHM values computed to be 288.26, 45.47 and 15.05 seconds for corresponding stimulus slopes of 0.01, 0.1 and 1°C/*s*, respectively. A similar trend was observed in the cGMP response dynamics with FWHM values decreasing from 268.42 seconds to 29.575 and 4.677 seconds as the stimulus steepness increased (shown in Fig. S5a-c).

**Figure 4.**
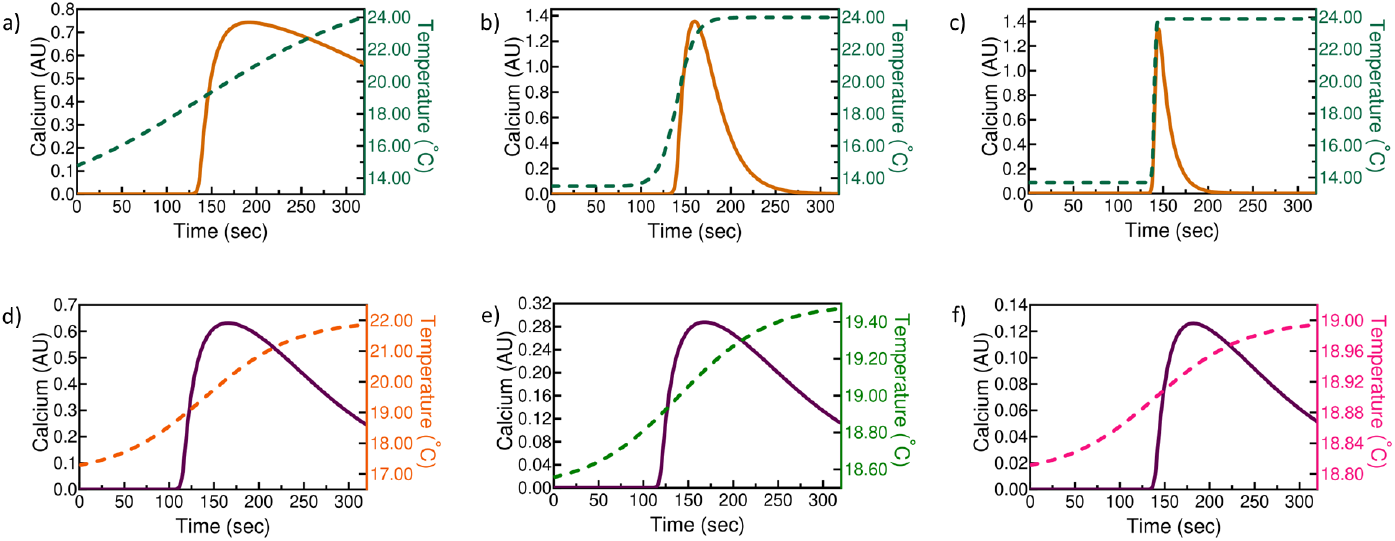
Effect of rate of change of signal on the calcium response: Spatial thermal signal with slope of **a)** 0.01°C/*s*, **b)** 0.1°C/*s*, **c)** 1°C/*s*. **Effect of temperature signal range on the calcium levels :** Range of the temperature signal is **d)** 5°C, **e)** 1°C, **f)** 0.2°C. The analysis of the model is performed at *T*_*C*_ = 20°C and the parameters used for signal, *S*, in Eq. 2 are listed in Table S2.

In addition, our analysis indicates a strong correlation between the rate of change of the signal and neural adaptation in the AFD neurons. Neural adaptation, which refers to the phenomenon of reduced neuronal activity in response to prolonged stimulation [53], is evident here. As seen in Fig. 4a-c, a faster thermal ramp (signal change) has a significant effect on response activity in terms of reduction in the total response duration, thereby driving it to steady-state values more rapidly. Further, on examining the dynamics of AFD neural adaptation, we find that the decay in the neuron’s response follows a single exponential adaptation across a broad range of stimulus steepness (shown in Fig. S6). The exponent fit on the decay of Ca^2+^ levels reveals a decrease in the relaxation times observed as 263.60, 31.03, 13.60 seconds, along with half-maximum decay times of 433.56, 187.87 and 154.96 seconds for stimulus slopes of 0.01, 0.1 and 1°C/*s*, respectively. Similar trends are observed for the cGMP response dynamics and the corresponding values are provided in Fig. S6a-c.

Although the rate of change in the stimulus increases 10 fold as we go from Fig. 4a-c, the neuron’s response dynamics (for both Ca^2+^ and cGMP) does not scale linearly with the rate of the stimulus. This non-linear relation is such that as the stimulus steepness increases, the relaxation time and FWHM of the AFD neuron’s response decrease at a faster rate. This correlation indicates that AFD neurons adapt more quickly and effectively to a faster changing stimuli and adjust their response dynamics accordingly.

Next, we investigate how effectively our model captures the dynamic range and sensitivity of the neuronal response. Experimental studies in the past have shown that AFD neurons are highly sensitive to temperature changes and can detect fluctuations even as small as 0.01°C/*s* across a wide range of signal magnitudes [10, 11, 58]. This precise sensitivity allows the worms to maintain isothermal tracking within a thermal deviation of 0.1°C for several minutes [10, 54]. In general, the transformation between temperature signal input and AFD neuronal response is characterized by measuring Ca^2+^ levels in response to temperature steps that vary across different ranges. This approach allows for quantifying how small temperature changes influence the activity of the AFD neuron. Here, we use our model to examine how neurons respond to decreasing temperature signal ranges of 5, 1, and 0.2°C (details in Table S2). The change in cGMP and Ca^2+^ levels are shown in Fig. S5d-f and Fig. 4d-f, respectively, for the temperature steps with varying ranges and a fixed slope of 0.02°C /*s*. Our model results show the adaptation response of the neuron to the signal for a step range as large as 5°C and as small as 0.02°C in a time period of about 320 seconds. Since the slope remains constant in Fig. 4d-f, the relaxation times during the decay of neuronal activity are nearly identical. However, the magnitude of the peak cGMP and Ca^2+^ levels attained is significantly affected by the range of the temperature step. These results from the model for varying slope and range of the temperature stimuli further advance our understanding of AFD’s neuronal response and its adaptation.

### Validation of model derived response dynamics of the AFD neuron

To assess the ability of our model in replicating the experimentally observed dynamics of AFD neurons, we compare our results against calcium imaging based response dynamics data reported by Huang *et al*. [28]. We do this to validate that our model, initially developed in relation to the experimental findings by Kobayashi *et al*. [27], translates well to another independent study as well. Signal-response data reported by Huang *et al*. shows a change in the Ca^2+^ levels in response to thermal stimuli varying in the range of 16-21°C. This calcium dynamics data recorded for the AFD soma in day one adult worms provides a basis for validating our model. Further, to ensure consistency between the experimental setup and our model, we adjust the thermal stimulus signal *S* accordingly (details mentioned in SI) and after introducing scaling model parameters, we find that the model accurately reproduces the Ca^2+^ dynamics observed in the experimental study. The comparison between the calcium dynamics reported in the experimental study and predicted by our model is shown in Fig. 5a.

**Figure 5.**
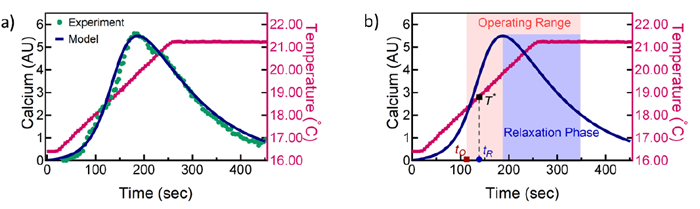
**a)** Comparison of calcium dynamics observed from the model (details in SI) to experimental study performed by Huang *et al*. [28]. **b)** Response properties of AFD neurons predicted from the the calcium level changes obtained from the model analysis.The thermal signal ranges between 16-21°C at a slope of 0.018°C/*s* and *T*_*C*_ = 20°C. *T* ^*^ and *t*_*R*_ are response temperature and corresponding time of response, respectively.

Further, we explore the response properties of the AFD neuron, which can be computed from the model’s predicted results and are commonly used to understand the effects of various conditions on the calcium response, as in previous experimental studies [27, 28]. Specifically, we examine three different properties: onset time of activation, operating range and relaxation phase of the AFD neuron’s response. These properties for the worms with *T*_*C*_ = 20°C and a thermal warming signal ranging from 16-21°C at a slope of 0.018°C/*s* are shown in Fig. 5b. The onset time of activation, *t*_*O*_, shown here is the time required for Ca^2+^ levels to reach a value equal to the maximum value divided by *e, i*.*e*., *C*_*max*_*/e*, starting from from *t* = 0, where *e* is the Napier’s constant. Similarly, the relaxation phase (blue region) is the time period required for maximal calcium levels (*C*_*max*_) to decrease till the value equal to *C*_*max*_*/e* on the decay side. The operating range of the AFD neuron (highlighted in pink) refers to the time interval between the onset time and the end of the relaxation phase of Ca^2+^ response levels. The model’s ability to reliably reproduce and validate key properties, as well as the calcium dynamics observed in experiments, enhances its credibility to study the intricate details of the AFD neuron’s response to thermal stimuli.

### High growth temperatures lead to delayed and reduced AFD neuron activity

After validating the model (Eqs. 15 and 12) through comparison with experimental data, we further extend its applicability and predict the response properties of AFD neurons for another range of thermal warming signal under different growth temperatures, *T*_*C*_. This demonstrates the model’s flexibility in adapting to varying thermal inputs within the optimal growth range for *C. elegans*, approximately between 15–25°C. While the model allows for visualization of the AFD neuronal response by analyzing calcium levels for worms conditioned across this entire range of *T*_*C*_, we focus on comparing the *T*_*C*_ evoked effects on response properties by presenting an analysis at three specific temperatures: 17, 20 and 23°C, shown in Fig. 6.

**Figure 6.**
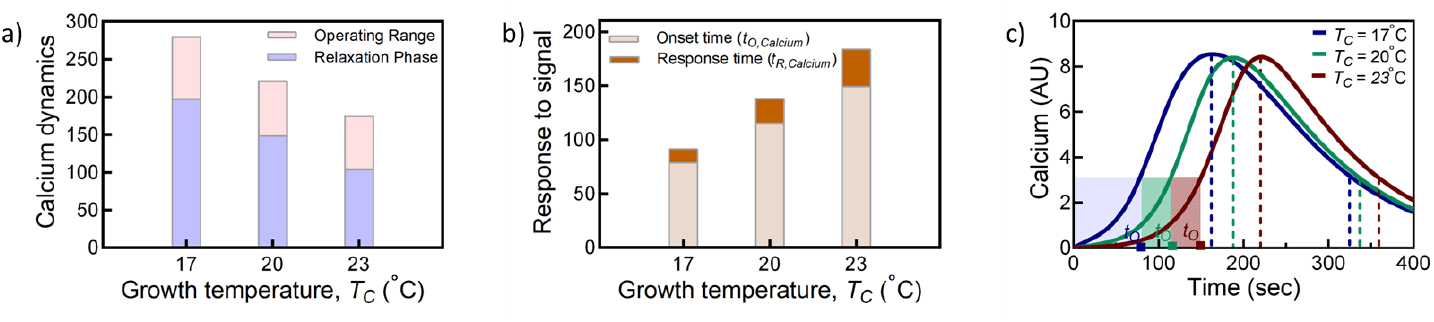
Effect of growth temperature on the response properties of AFD neuron analyzed from the change in calcium levels: Comparison of **a)** the duration of operating range and relaxation phase, **b)** onset time of activation, *t*_*O*_, and response time, *t*_*R*_, **c)** calcium dynamics as response of AFD neurons for worms with *T*_*C*_ = 17, 20 and 23°C.

Upon linear thermal warming from 14-24°C, the operating range and relaxation phase for worms with *T*_*C*_ = 17°C are 280 and 197 seconds, respectively. For an increase in *T*_*C*_ to 20°C, these durations decrease to 221 and 149 seconds and a further decrease occurs at *T*_*C*_ = 23°C where the values are found to be 175 and 104 seconds. This indicates that higher *T*_*C*_ values result in a reduced operating range and relaxation phase of the response elicited by AFD neurons (see Fig. 6a and c), thus narrowing the FWHM of the calcium dynamics, as shown in Fig. S7. In contrast, the values for onset time of activation, *t*_*O*_ and time of response, *t*_*R*_, increase with increasing *T*_*C*_ (see Fig. 6b-c). This suggests an overall delay in the response of neurons, characterized by both a delayed onset of activation as well as temporal response to the thermal stimuli at higher *T*_*C*_. Additionally, our model reveals that the growth of worms at elevated temperatures results in an increased difference between *t*_*O*_ and *t*_*R*_. This difference changes from 12.7 seconds at *T*_*C*_ = 17°C to 22.8 seconds at *T*_*C*_ = 20°C and then to 34.4 seconds at *T*_*C*_ = 23°C. These observations further indicate that the ability of worms to exhibit a fast response to temperature changes decreases when they are grown at high temperatures.

## SUMMARY AND DISCUSSION

Temperature in an organism’s environment is a fundamental sensory cue that impacts survival and complex social behaviors. The ability of organisms to respond to temperature variations is also crucial for their adaptation to changing conditions. Over the years, *C. elegans* has been widely used as a model system to explore the fundamental mechanisms of thermosensation and the temperature-dependent behaviors [1]. These tiny worms are known to detect environmental temperature mainly through a pair of head sensory neurons, the AFD neurons, which play a major role in regulating the thermotactic behavior of *C. elegans* [3, 10, 26, 59]. The experimental research on thermosensation in worms has focused extensively on examining the response of AFD neurons under changing temperatures where several key studies have also shed a light on their functional role [2, 3, 10, 11, 26, 34]. However, the mathematical understanding of the signal response mechanism in these neurons has received considerably less attention [17, 18, 60]. Developing theoretical models to understand how AFD neurons process external thermal information and generate a response is important to uncover the underlying neuronal mechanism. A mathematical framework can reveal how the AFD neuron modulates calcium dynamics in response to varying thermal stimuli, offering deeper insights into signal processing pathways. Moreover, to fully capture the dynamic nature of the responses elicited by the AFD neuron, a robust model capable of providing quantitative predictions across a broad range of thermal inputs is necessary.

Here, in this work, we develop and analyze a mathematical model that decodes the intricate dynamics of thermosensation in *C. elegans*, focusing on the AFD thermosensory neurons. In the past, a functional model to understand themosensation by AFD neurons in worms has applied a linear-nonlinear convolution model [17]. This approach of analyzing the relationship between temperature and calcium activity in the neurons has been valuable in defining the response function and has significantly advanced network models of the full thermotaxis circuit. However, the linear-nonlinear model does not incorporate the underlying biophysical mechanisms believed to drive AFD thermosensation. Other theoretical works such as by Nakazato *et al*. [61] focus more on the effect of thermal signal on the migration behavior of worms where a biased random walk was used to model population dynamics of *C. elegans* across a thermal gradient. Another type of model studying AFD neurons rely on Hodgkin-Huxley formalism to develop a conductance based model that provides biophysical representation of the neurons [60]. As an alternative to these models, another approach to studying neuronal response is by constructing a phenomenological model, that accurately represents the biochemical processes underlying the sensory mechanism of the neuron. In this work, we adopt this approach primarily because phenomenological models tend to generalize more effectively and are instrumental in making quantitative predictions about the system’s response to stimuli.

Our model on the study of thermosensory response of AFD neurons in *C. elegans* integrate the known biochemical reactions of the main signaling pathway, *i*.*e*., the cGMP signaling cascade, in the neurons. The model provides a quantitative framework for the pathway illustrated in Fig. 1, with an aim to understand the dynamic processes of signal transduction in AFD neurons during linear thermal warming. The analysis mainly focuses on the dynamics of two key components-cGMP and calcium - described by Eqs. 11 and 12, which play a central role in modulating the neuronal activity as well as signal transmission. The primary analysis of the model is performed for a thermal ramp signal varying within the physiological range of *C. elegans* (∼15°C - 25°C), in line with previous experimental studies on AFD neuron responses. First, the model analysis shows that both cGMP and calcium responses in AFD neurons adapt to the first derivative of the temperature stimulus, rather than to the absolute levels of temperature (shown in Fig. 2). This allows AFD neurons to detect temperature changes more sensitively, enabling the worm to respond even to subtle thermal shifts. The model integrates the concept of temperature memory in Eqs. 14 and 15, showing how AFD neurons retain and reset their responses based on the growth temperature, *T*_*C*_, of the worms. This memory affects the neuron’s response to temperature stimuli, with the calcium and cGMP responses initiating at a threshold temperature near *T*_*C*_. This suggests that the thermosensory system of the worms is calibrated to their developmental conditions and is shown in Fig. 3a-c, a finding that has been corroborated by previous experimental studies [26, 27]. The model offers the advantage of examining the relationship between growth temperature conditions and the neuronal response threshold across a full range of stimuli, a relationship that is challenging to study experimentally.

In this study, we further elucidate how the AFD neuron’s response is influenced by rate and range of temperature signals, where we find that fast warming leads to fast increase in calcium levels, thus attaining peak values within short time periods of signal stimulation. We find that rapid rate of input signal also results in faster neural adaptation, thus leading to faster exponential decays and correspondingly, low relaxation times for the response dynamics of cGMP and calcium. Notably, the model is also sensitive and effectively captures the response of the neuron over a wide range of temperature steps (0.2 °C - 5°C). This emphasizes the intricate relationship between the temporal change of the signal to the response dynamics elicited in the AFD neuron. Next, the model is validated by comparing its predictions to experimental calcium dynamics reported by Huang *et al*. [28] for worms exposed to thermal stimuli between 16-21°C. The model results detailed in Fig. 5 shows a very good match to the experimentally observed calcium responses. Further, through the model analysis (shown in Fig. 6), we find that worms cultivated at higher temperatures (23°C) exhibit delayed and shortened calcium responses compared to worms grown at lower temperatures (17°C). This suggests that higher growth temperatures lead to a narrowed operating range and faster adaptation, reflecting a quantitative viewpoint about how organisms adjust its sensitivity based on its past environmental experience.

Overall, this study introduces a model that effectively captures the signal response mechanism of AFD neurons in *C. elegans*. By utilizing a minimal number of parameters and a straightforward set of equations, the model provides an efficient and effective framework for understanding the core response dynamics of thermosensation. With a focus on the main components of the signaling pathway, we reduce complexity without sacrificing accuracy and make the model easier to interpret, modify and apply. Moreover, the simplicity and reliance of the model on the fundamental principles of neuron sensation such as intracellular signaling, calcium dynamics and sensory adaptation broadens its applicability to understand other thermosensory systems. The mechanisms modeled here, like second messenger signaling and temperature dependent calcium influx are prevalent across many organisms from invertebrates to mammals. Despite its reduced complexity, the model successfully replicates key experimental observations for the AFD neurons such as temperature-evoked memory effects and neural adaptation. This makes it a powerful tool for theoretical investigations and a guide for future experimental work. Ultimately, this approach not only captures key mechanisms of thermosensation in the *C. elegans* model system, but also offers a valuable framework for refining our current understanding of sensory processes across various biological contexts.

## MATERIALS AND METHODS

### Signal in the model to validate against experimental data

The ‘curve fit’ function from SciPy’s ‘optimize’ module was used to perform non-linear least squares fitting of sigmoidal function (Eq. 2) to the signal data used in experimental studies by Kobayashi *et al*. [27] in Fig. 1D and by Huang *et al*. [28] in Fig. 3g, respectively. The model function parameters were optimized using the ‘dogbox’ method, a trust-region algorithm suitable for non-linear problems with well-behaved models. After providing the initial parameter guesses, the optimized parameters and their covariance matrix were obtained, with the optimal parameter values representing the best fit of the sigmoidal function (signal) to the data.

### Computational Methods

All numerical integration to solve Eqs. 11 and 12 was performed with Mathematica’s [62] NDSolve function by selecting automatic method which allows it to choose the most appropriate solver based on the system’s characteristic. The accuracy and precision goals were set high enough to ensure reasonable numerical convergence.

### Relation between response time and growth temperature

To establish the relationship between the response temperature, *T* ^*^, and the growth temperature, *T*_*C*_, the experimental data from Kobayashi *et al*. [27] is used. The corresponding values of *T*_*C*_, *T* ^*^ and the response time, *t*_*R*_ from [27] are provided in Table S1. A linear fit to the relationship between *T*_*C*_ and *T* ^*^ in Table S1 yields the equation:

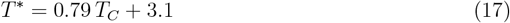

To determine the corresponding response time, *t*_*R*_, the absolute value of *T* ^*^ for a given *T*_*C*_ (determined from Eq. 17) is used in the following equation:

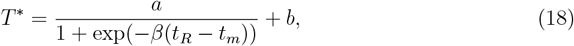

where the parameter *a* scales the range of the temperature signal and the constant *b* sets the offset temperature in the function. The parameter *β* determines the rate of transition between temperatures *b* and *a* + *b*, while *t*_*m*_ is the characteristic time point that represents the midpoint of the sigmoidal transition.

From Eq. 18, the respective values of response time, *t*_*R*_, obtained from our model for *T*_*C*_ values of 17, 20 and 23 °C are 95.5, 134.5 and 186.7 seconds, respectively. This relationship for *T*_*C*_ values varying between 16 to 24°C is also plotted in Fig. S4a.

### Experimental data used in this study

To compare the model predictions, response temperature values (*T* ^*^) from Kobayashi *et al*. [27], shown in Fig. 3e of our study, were extracted from Fig. 1E (lower panel) in [27]. Similarly, for validation against experimental data from Huang *et al*. [28], presented in Fig. 5a of our study, data were obtained from Fig. 3g in [28]. All data extraction was carried out using the WebPlotDigitizer software [63].

## Data, Materials, and Software Availability

All study data are included in the article and/or supporting information. Code and data used for the results in this paper can be found at https://github.com/cebpLab/AFD-Model.

## Notes

https://github.com/cebpLab/AFD-Model

### Competing Interest Statement

The authors have declared no competing interest.

