## Supporting Information for "Neuronal Decoding of Temperature Signals in *Caenorhabditis elegans*"

**This PDF file includes:**

Supporting text

Figures S1-S7

Table S1-S2

SI References

[2, 3],

### SUPPORTING INFORMATION

#### Calculation of $k_r$

The numerical value of the rate constant for the production of cGMP, $k_r$, in Eq. 4 of the main text is estimated from the data reported for the change in the guanylyl cyclase activity of GCY-12 as a function of temperature in the study by Yu *et al.* [1]. The specific activity of the guanylyl cyclase receptor is defined as the moles of cGMP formed by GCY-12 in a given amount of time divided by total protein mass in the reaction volume. This definition can be expressed as,

$$\text{Specific activity} = \text{Rate of reaction} \times \frac{\text{Volume of reaction}}{\text{Mass of total protein}} \tag{S1}$$

As stated in the study by Yu *et al.* [1], the volume of reaction $= 100\mu L$ and the mass of membrane proteins is $\sim 35\mu g$. Therefore, Eq. S1 with units for the specific activity taken in $pmol\, mg^{-1}\, min^{-1}$ can be written as

$$\begin{aligned} \text{Rate of reaction} &= 0.35 \times \text{Specific activity} \left( \frac{pmol\, mg^{-1}\, min^{-1}\, \mu g}{\mu L} \right) \\ &= 0.35 \times 10^{-15} \times \text{Specific activity} \left( mol\, min^{-1}\, \mu^{-1} L^{-1} \right) \\ &= 0.35 \times 10^{-9} \times \text{Specific activity} \left( M\, min^{-1} \right) \\ &= 0.00583 \times 10^{-9} \times \text{Specific activity} \left( M\, s^{-1} \right) \end{aligned} \tag{S2}$$

As for a first order enzymatic reaction, the rate of reaction of cGMP formation can be given as $k_r$[GTP]. Considering [GTP] $= 100\, \mu M$ [1], $k_r$ is given by

$$k_r = 583 \times 10^{-10} \times \text{Specific activity} \left( s^{-1} \right) \tag{S3}$$

The specific activity for GCY-12 are 39, 62 and 67 $pmol/mg/min$ at 15, 20, 25°C, respectively [1]. Therefore, taking an average of these three values shows that an estimated value of $k_r$ will be $3.2648 \times 10^{-6} s^{-1}$.

#### Calculation of $E_f$

The constant $E_f$ in Eq. 8 of the main text changes the electrical current unit to concentration changes while taking into account the volume of the AFD neuron and is defined as [2, 3],

$$E_f = \frac{f_{\mathrm{Ca}^{2+}}}{2\mathcal{F} v_{AFD}} \tag{S4}$$

where,
$f_{Ca^{2+}}$ is the fraction of current carried by $Ca^{2+}$ ions and has value 0.2 (from [3]),
$\mathcal{F}$ is Faraday's constant with value 96485 $Cmol^{-1}$,
$v_{AFD}$ is the average volume for the pair of AFD neurons with value 32.055 $\mu m^3$ (as given in https://neuromorpho.org/). Therefore,

$$E_f = \frac{0.2}{2 \times 96485 \times 32.055\, C\, mol^{-1}\, \mu m^3} = 32.33\, \mu M C^{-1} \tag{S5}$$

#### Normalization of cGMP and calcium concentrations

The cGMP and $Ca^{2+}$ concentrations obtained from the mathematical model analysis are normalized to their baseline values to depict the response in arbitrary units relative to the temperature stimulus. The relation between cGMP and $Ca^{2+}$ concentrations and their respective response is analyzed as,

$$\text{cGMP response} = \frac{\text{cGMP concentration } (\mu M)}{10^{-4}(\mu M)} \tag{S6}$$

$$\text{Ca}^{2+} \text{ response} = \frac{\text{Ca}^{2+} \text{ concentration } (\mu M) - 10^{-1}(\mu M)}{10^{-1}(\mu M)} \tag{S7}$$

#### Model validation to the experimental data

For the validation of model to the experimental data for $Ca^{2+}$ levels shown for day one worms grown at $T_C$ = 20 °C by Huang *et al.* [4], the signal, $S$, in Eq. 2 is modified with values of parameters as $a$=5.35°C, $b$=16°C , $\beta$=0.018$s^{-1}$, $t_m$=129$s$.

The parameter values are set as $\tau = 16.6s^{-1}$ and $t_R = 138.37s$ in Eq. 14 of the main text for the activated receptor $R_a$ with memory recall function, and are optimized to fit the model to experimental data in [4]. To quantitatively compare the model results with the experimental data, the $Ca^{2+}$ response levels are scaled by a numerical factor of 10. All other parameter values remain consistent with those listed in Table 1, and the comparison is presented in Fig. 5a of the main text.

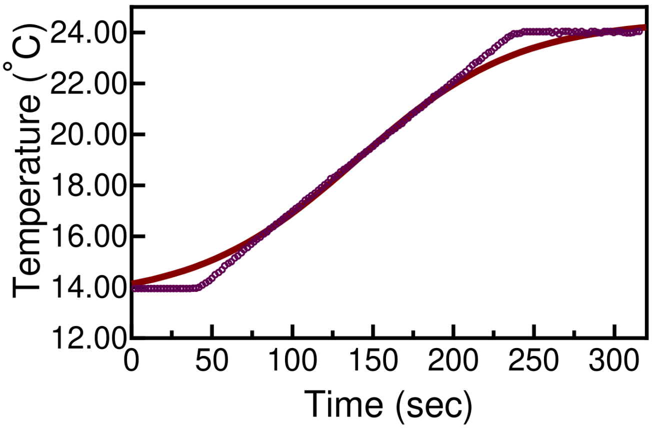


Figure S1. **Stimulus in the model:** is represented by a sigmoidal function (Eq. 2) with numerical values of parameters $a = 11°\text{C}$, $b = 13.5°\text{C}$, $\beta = 0.02s^{-1}$, $t_m = 140s$ to imitate the linear thermal warming signal employed in experimental studies [5] on the AFD neurons.

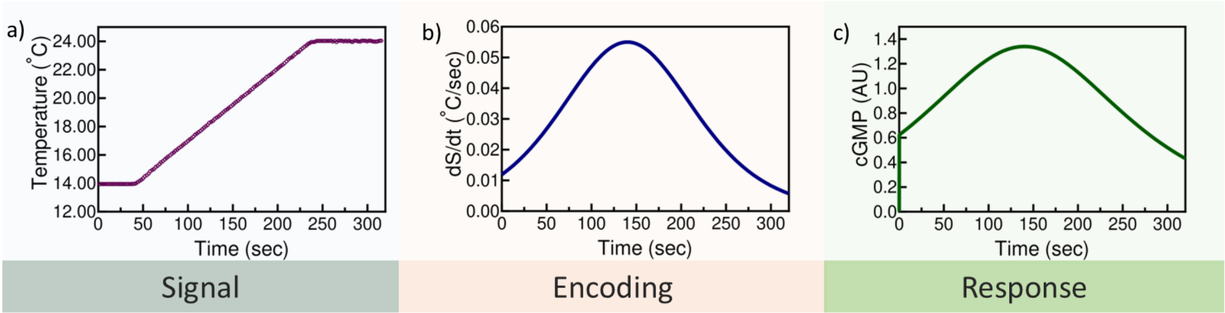


Figure S2. **Signal processing in AFD neurons: a)** Linear thermal warming signal employed in experimental study [5]. **b)** Encoding of signal as first derivative, $dS/dt$, (Eq. 13) by the neurons. **c)** cGMP levels observed through the model analysis for signal encoded as $dS/dt$ at $T_C = 20°$C.

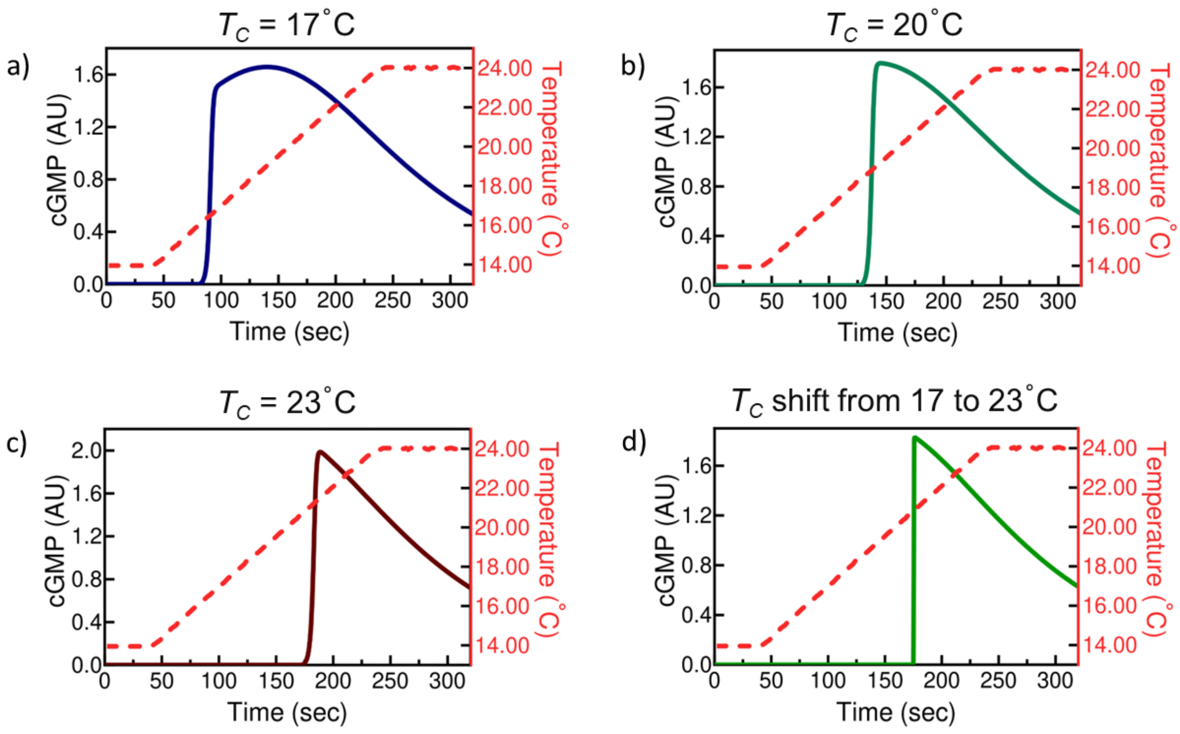


Figure S3. **Effect of growth-temperature memory on cGMP levels:** cGMP response dynamics observed through the model analysis in AFD neurons for linear thermal ramp as signal (Eq. 2 and Fig. S1) at **a)** $T_C = 17°$C, **b)** $T_C = 20°$C, **c)** $T_C = 23°$C, **d)** Shift in $T_C$ from 17°C to 23°C.

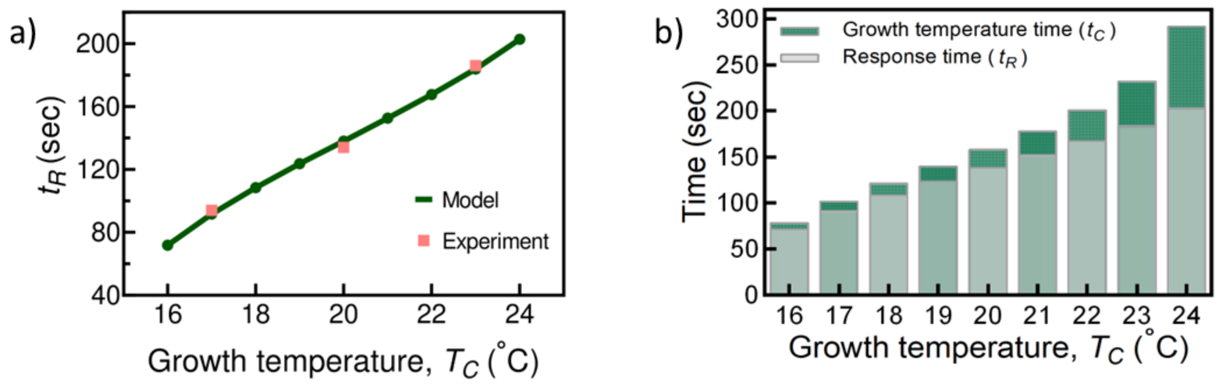


Figure S4. **a)** Comparison between response time, $t_R$, of the neurons obtained from the model to the experimental values observed in [5], **b)** Change in the value of response time, $t_R$, for calcium levels at time, $t_C$ (which is corresponding time to reach the growth temperature, $T_C$), for linearly increasing temperature stimulus in the range of 16°C to 24°C.

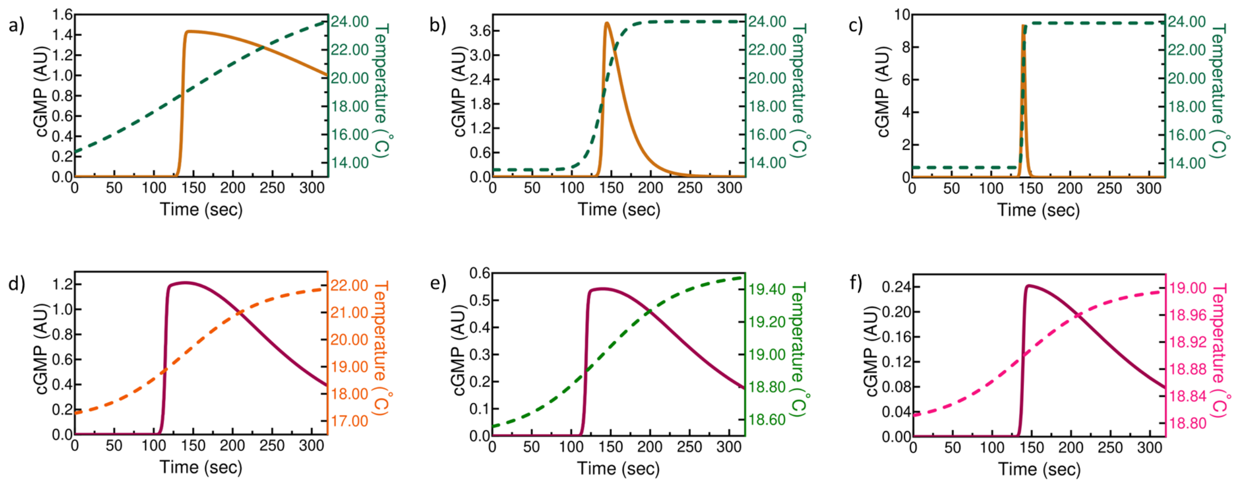


Figure S5. **Effect of rate of change of the signal on the cGMP response:** Spatial thermal signal with slope of **a)** 0.01°C/$s$, **b)** 0.1°C/$s$, **c)** 1°C/$s$. **Effect of temperature signal range on the cGMP levels:** Range of the temperature signal is **d)** 5°C, **e)** 1°C, **f)** 0.2°C. The analysis of the model is performed at $T_C = 20$°C and the parameters used for signal, $S$, in Eq. 2 are listed in Table .

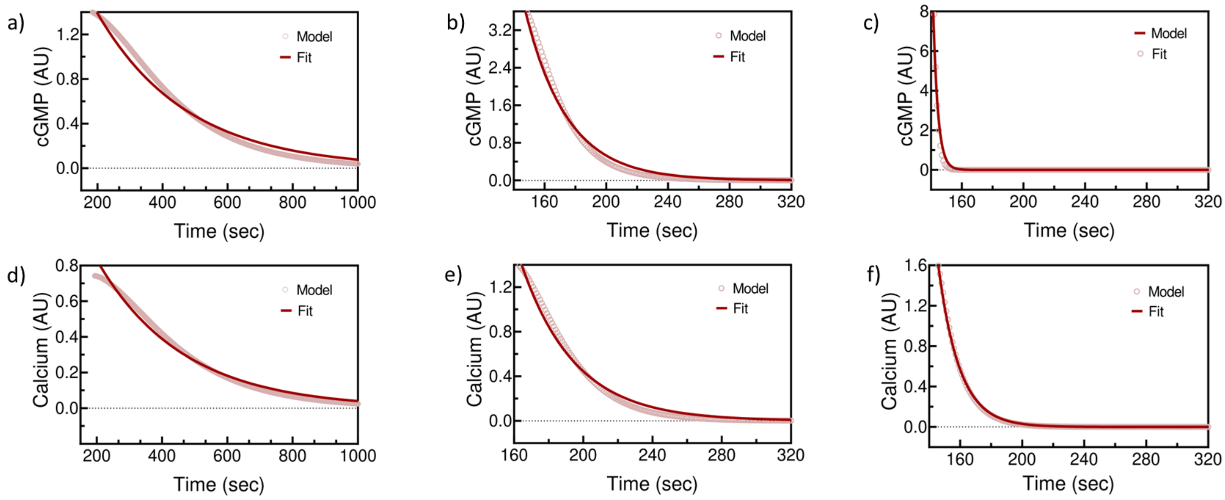


Figure S6. **Dynamics of neural adaptation:** Fitting of single exponent function $a\,Exp(-\frac{1}{\tau}(t-t_0))$ to the decay of cGMP levels **a)** in Fig. S5a with $a = 1.39$, $\tau = 276.03$, $t_0 = 200$, **b)** in Fig. S5b with $a = 2.28$, $\tau = 26.23$, $t_0 = 160$, **c)** in Fig. S5c with $a = 12.6841$, $\tau = 2.7342$, $t_0 = 140$. Fitting of single exponent $a\,Exp(-\frac{1}{\tau}(t-t_0))$ to the decay of calcium levels **d)** in Fig. 4a with $a = 0.83$, $\tau = 263.60$, $t_0 = 200$, **e)** in Fig. 4b with $a = 1.61$, $\tau = 31.03$, $t_0 = 160$, **f)** in Fig. 4c with $a = 2.46742$, $\tau = 13.60$, $t_0 = 140$.

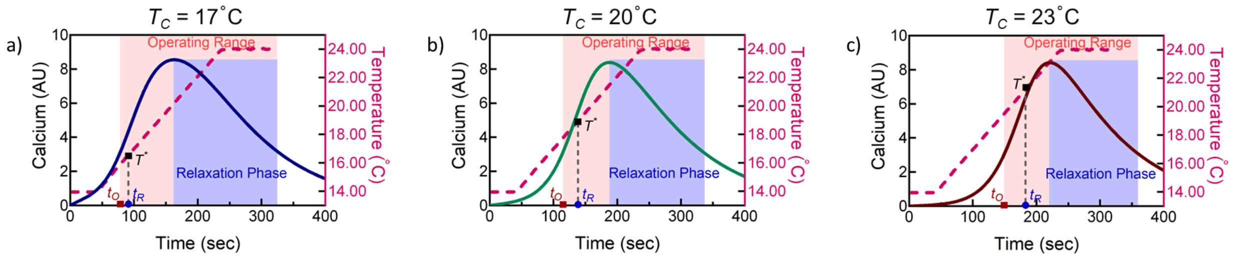


Figure S7. **Effect of growth temperature on the response properties of AFD neuron analyzed from the change in calcium levels at: a)** $T_C = 17°\text{C}$, **b)** $T_C = 20°\text{C}$, **c)** $T_C = 23°\text{C}$.

| $T_C$ (°C) | $T^*$ (°C) | $t_R$ ($s$) |
|---|---|---|
| 17 | 16.7 | 94 |
| 20 | 18.7 | 134 |
| 23 | 21.4 | 186 |

Table S1. Experimental data (Fig. 1E, lower panel) from Kobayashi *et al.* [5] used to observe the relationship between growth temperature ($T_C$), response temperature ($T^*$) and response time ($t_R$).

| Figure No. | $a$ (°C) | $b$ (°C) | $\beta$ ($s^{-1}$) |
|---|---|---|---|
| 4a, S5a | 14 | 12 | 0.01 |
| 4b, S5b | 10.5 | 13.5 | 0.1 |
| 4c, S5c | 10.2 | 13.7 | 1 |
| 4d, S5d | 5 | 17 | 0.02 |
| 4e, S5e | 1 | 18.5 | 0.02 |
| 4f, S5f | 0.2 | 18.8 | 0.02 |

Table S2. The mathematical expressions for signal, $S$, in Eq. 2 used to perform the model analysis described in Fig. 4 and Fig. S5 with $t_m$=140$s$.
